# The Connectome Interpreter Toolkit

**DOI:** 10.1101/2025.09.29.679410

**Authors:** Yijie Yin, Judith Hoeller, Alexander Mathiasen, J. M. F. Tsang, Monique Estelle Charrier, Albert Cardona

## Abstract

Complete synaptic wiring diagrams, or connectomes, of whole brains open new opportunities for studying the structure-function relationship of neural circuits. However, the large number of nodes and edges in the graphs makes the analysis challenging. Here, we present the **Connectome Interpreter**, an open-source software toolkit for efficient graph exploration to find polysynaptic pathways, compute the effective connectivity and receptive fields for arbitrarily deep neurons, slice out subcircuits, and construct non-linear but differentiable circuit models, implemented using efficient approaches tailored to large connectomes with abundant divergent and convergent connections, such as that of the fruit fly *Drosophila melanogaster*. Our approach delivers results orders of magnitude faster than conventional methods on consumer hardware. We demonstrate the capabilities of our toolkit with select applications, including quantifying the density of polysynaptic connections in the whole adult fruit fly brain, exploring the necessity for non-linearities in circuit modeling, and combining known function of neurons with the connectome to aid in formulating hypotheses of circuit function.

## 1 Introduction

Neural circuit architecture constrains the functional capabilities of brains, with mapped neuronal wiring diagrams, or connectomes, serving as a basis for disambiguating competing hypotheses of neural function [1]. Recent large-scale research projects densely reconstructed neurons and labeled their synapses from nanometer-resolution volumes of the brains of organisms ranging from the fruit flies to human [2–15]. These large, directed, weighted, signed (excitation/inhibition) connectomes establish a strong foundation for studying brain function: all circuits are now accessible for analysis.

However, connectomic datasets reveal unexpected complexity: an average neuron connects to ∼ 130 other neurons (median ∼ 90) and ∼ 80 cell types (median ∼ 55) in the central brain of the fly, indicating that the synaptic connectivity graph is dense enough to render circuit analysis a challenge. While a common approach is to focus on the stronger synaptic weights, the continuous distribution of connection weights makes defining a well-justified threshold challenging. Moreover, because most connections are weak, thresholding effectively removes most of the connections, inadvertently discarding potentially crucial signal integration patterns.

The observed dense synaptic connectivity of brain connectomes indicates that many neurons simultaneously participate in multiple sensory-motor pathways underlying natural behaviors. Yet, knowledge of neuronal function typically advances by experimentally manipulating and monitoring specific neurons with controlled, limited stimuli. Such an approach poses a fundamental scaling problem: exhaustively testing the relationships among combinations of sensory inputs, neuronal activations, and behavioral outputs is infeasible.

The study of the relationship between circuit structure and function fundamentally involves linking the known to the unknown (Figure 1A) — a central theme of neuroscience since its inception. Whether mapping sensory receptive fields or assigning functional roles based on circuit interactions, neuroscientists leverage prior data (e.g., characterized sensory neurons or behaviors) to interpret unexamined neural circuits. The advent of multiple connectomic datasets amplify both the need and opportunity for new methods that systematically bring functional data into circuit analysis.

**Figure 1:**
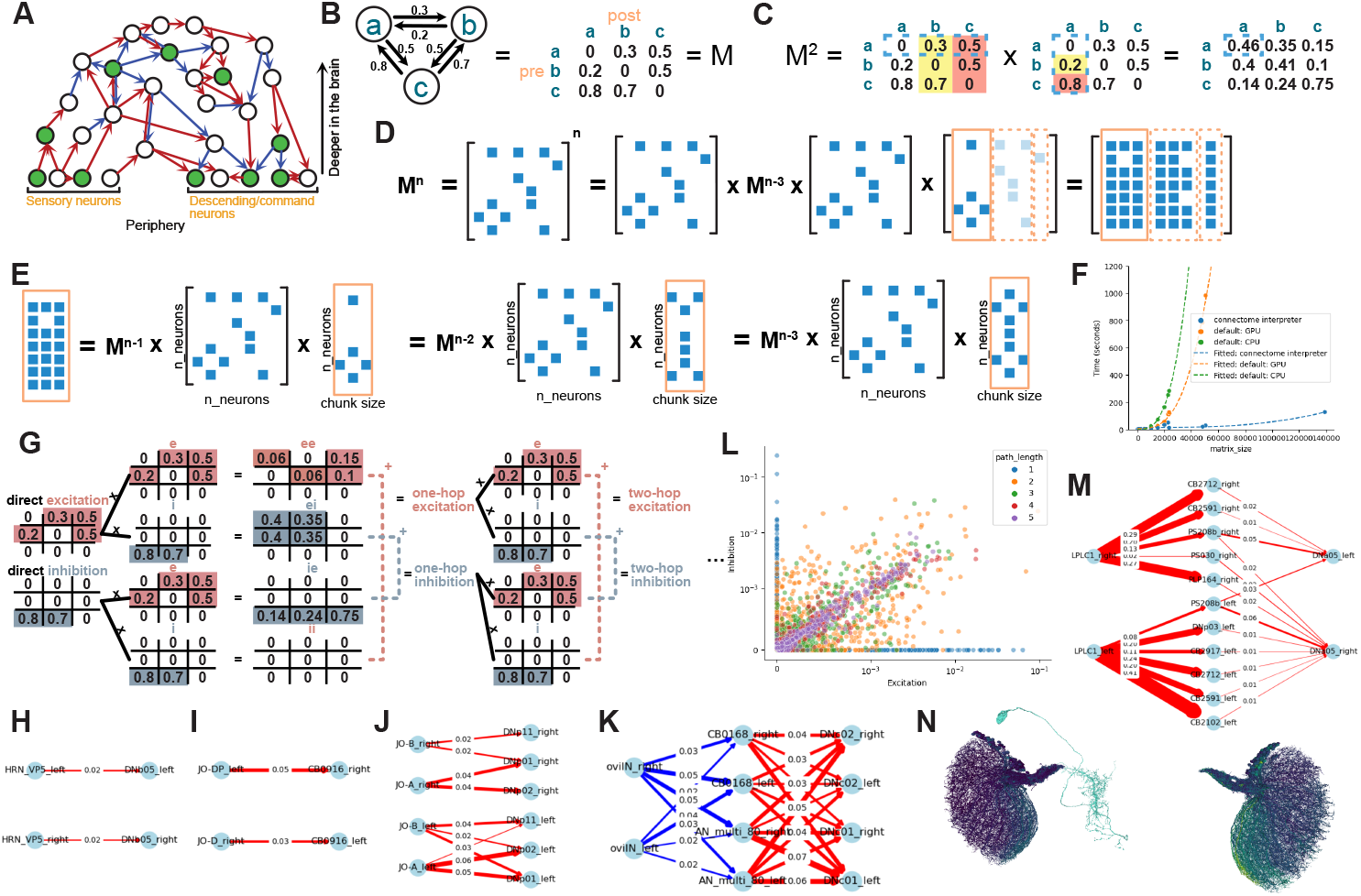
Linear approaches of target neuron selection with examples. **A**. Illustration of connectome-based hypothesis generation. Green fill indicates existing research on that neuron. **B**. Illustration of the equivalence of two representations of a network M. **C**. Calculation of all-to-all two-hop effective connectivity is equivalent to multiplying the connectivity matrix with itself once. The blue dashed boxes highlight the numbers multiplied to yield 0.46, and the fill colors show the “column picture of matrix multiplication”: in this case, the first column of the result matrix is the sum of: 0 × the first column, 0.2 × the second column, and 0.8 × the last column. **D-E**. We combine sparsity with chunking, by dividing the last matrix into one dense column chunk at a time, and multiplying it with all the sparse matrices before (E), resulting in a dense matrix the size of the chunk. We then combine the (thresholded) dense column chunks together. **F**. Comparison of our sparse chunked method with dense matrix multiplication on GPU and CPU, for different connectivity matrix sizes. **G**. To calculate signed effective connectivity, we divide the connectivity matrix into excitatory rows and inhibitory rows, and multiply each separately with excitatory and inhibitory rows, for all combinations of *ee, ei, ie*,ii. We then add *ee* and *ii* together for the “new” excitatory rows; and *ei* and *ie* together for the “new” inhibitory rows. **H**. *HRN V P* 5 connects to the ipsilateral *DNb*05. The edge labels indicate normalized input for *DNb*05. **I**. *JO* − *D* connects to the contralateral *CB*0916. **J**. *JO* − *A* and *JO* − *B* connect to the ipsilateral *DNp*01 (giant fiber), *DNp*02 and *DNp*11. **K**. oviLN connects bilaterally to *DNc*01 and *DNc*02 through indirect inhibition. **L**. Direct and effective signed connectivity between 100 randomly-selected cell types, colored by the path length. **M**. *LPLC*1 connects bilaterally, indirectly, to *DNa*05, through excitatory connections. The connections are stronger contralaterally. **N**. Neuroglancer scene snapshot, visualizing two-hop effective connectivity from individual *LPLC*1 neurons, each in its own color, to *DNa*05 (cyan). Yellow means stronger connectivity. All wiring diagrams are thresholded at 1% normalized input.

To address these challenges, we developed an approach to study neuronal function in which the connectome serves as not just a lookup table but an effective basis for translating structural complexity into functional insights.

Here, we introduce *Connectome Interpreter*, an open-source toolbox (https://github.com/YijieYin/connectome_interpreter) specifically developed to analyze circuit function on the basis of combining connectomes with known but potentially sparsely-annotated neuronal function. The toolbox accelerates common network analysis operations such as effective connectivity calculation [16–19] and polysynaptic path finding, while enabling exploration of previously inaccessible questions, including the identification of the stimulus optimal for activating neurons of interest. We demonstrate the utility of our toolbox through highlighting neural circuits of potential functional significance, and examining the role of non-linearity in signal propagation.

## 2 Results

### 2.1 Table of neurons with known functions

Knowledge of the function of at least some cell types is essential to infer the function of any particular cell type, by propagating functional information across the circuits comprised in a connectome. While connectomes are generally only now starting to be available, literature on the function of cell types has been accumulating for decades in multiple animal models.

We have collected information on the function of known cell types from the fly neuroscience literature into a table (https://tinyurl.com/known-neuron-function), with 562 cell types. Of these, 402 are from [11] (of which we extended further details on 61), 113 are from us, and the rest are from the community (see Acknowledgements).

The table shows that there is partial knowledge on at least 30% of the cells, comprising 5% of all cell types. In particular, the table includes 2.4% (174/7175) of the central brain cell types, 15% (71/481) of the descending cell types, 17% (60/352) of visual projection, and 4% (15/423) of the sensory neurons, indicating that more is known closer to the sensory/motor periphery of the central nervous system (as illustrated by Figure 1A).

We recognize that literature coverage of cell types continuously evolves, and therefore we opened up the table to the neuroscience community to further contribute by reviewing, correcting, and adding new entries via: https://tinyurl.com/known-neuron-function, to ensure all relevant work is represented. For maximal accessibility, we integrated the table into the Codex data curation initiative (https://codex.flywire.ai, 20), and include a snapshot in Supplementary table 1.

### 2.2 Connectome-driven target neuron selection

Our goal is to deliver effective tools for, given sparse functional data and dense connectomic data, identifying specific, understudied neurons for subsequent and targeted experimentation. Here, we provide both linear and non-linear approaches to tackle this analysis.

#### 2.2.1 Linear approach: effective connectivity

The effective connectivity approximates the amount of influence a source neuron could have on a target neuron, across all polysynaptic pathways of length *n*. One popular approach to estimating the effective connectivity is to multiply connection weights along each path from a source to a target neuron, and then add up this product across all parallel pathways between the same source and target [16–19].

However, multiplying and summing connection weights, which is equivalent to taking the power of the connectivity matrix (Figure 1B-C), is in practice limited to networks with a small number of nodes (a node in this case is a neuron in the connectome), because the computational cost of multiplying two *n* × *n* connectivity matrices scales to the third power of the number of nodes (*n*), and linearly with the number of hops (*h*) in the longest polysynaptic path to consider (*O*(*n*^3^ · *h*)) (Figure 1F, Methods 5.0.1).

Our method avoids the *n*^3^ scaling by first leveraging the sparsity of the fly brain connectome: for its 140,000 neurons, on average any one neuron receives synapses from 1/1,000 of all neurons, i.e., about 140, when considering all mapped synapses. The sparse matrix representation brings huge savings in computer memory usage and number of operations to perform, skipping all the zeros during matrix multiplication.

However, the divergence and convergence of the connectivity (Figure 4A) entail that the matrix products quickly become denser the more matrices are multiplied, rendering sparse representation no longer useful for longer polysynaptic hops. We address this challenge by dividing the dense products into chunks with a customizable size (Figure 1D-E), such that only one small dense chunk has to be held in memory at any one time, drastically reducing the memory requirement for such large-scale operations.

Our approach makes the same computation orders of magnitude faster, taking minutes instead of hours or days to compute the effective connectivity between all neurons (Figure 1F), for a connectome the size of the brain of the adult fly brain, at ∼ 140,000 [2, 3]. Further, we cater to a diversity of available resources in typical neuroscience labs, by making our implementation customizable in memory consumption, facilitating execution on widely-accessible computational resources, from a laptop to Google Colab, when conventional approaches (such as dense matrix multiplication; see Methods 5.0.1) otherwise require high performance computing resources for these datasets. The results can then be saved and used for instant queries of effective connectivity between any groups of neurons.

As a proof of principle, we combine the (effective) connectivity matrix with our table of functionally characterized neurons (Supplementary table 2 and 3). This yields, for every cell type, a “contextualized receptive field” (RFc), which we define as the receptive field in terms of not only peripheral sensory neurons (the ‘conventional’ receptive field), but also the functionally characterized neurons anywhere in the connectome. This approach was previously illustrated for the receptive field of the lobula plate tangential cells (LPTCs) via their direct T4/T5 inputs [21]. To illustrate the utility of the contextualized receptive field, we look for unreported (direct and indirect) putative functional relationships between reported cell types in the literature.

We consider the following monosynaptic (direct) connections:

1. [22] showed that *Drosophila melanogaster* prefer humidity. Correspondingly, we find that the HP5 glomerulus of the hygrosensory receptor neurons, sensitive to humidity [23], synapse onto the ipsilateral side of DNb05 which is responsible for ipsiversive turning (turning towards the ipsilateral side; [24]) (Figure 1H). Through effectively combining the existing knowledge of cell types with the connectivity matrix, we identify the HP5-to-DNb05 connection as potentially the neural circuit basis for turning towards the side with higher humidity, a hypothesis that could now be tested experimentally.
2. Johnston’s organ neurons, JO-D, shown to signal static deflection of the antenna [25], are connected to the contralateral CB0916 which is involved in neck motor control [26] (Figure 1I). The direct synapse from JO-D to CB0916 could therefore drive neck movement upon antennal deflection.
3. Johnston’s organ neurons JO-A and JO-B detect sound [27, 28]. We find that they synapse directly not only to the Giant Fiber neuron as shown before [29], but also onto additional descending neurons (DNp02 and DNp11) implicated in directional escaping in response to visual input from different angles [30] (Figure 1J). We thus hypothesize that these descending neurons could also drive directional escaping in response to sound.

We considered as well the following polysynaptic (indirect) connection: [31] showed that oviINs inhibit oviDNs responsible for oviposition. We show that they also indirectly inhibit DNc01 and DNc02 (Figure 1K), two cell types known to release SIFamide [32, 33]. Activation of SIFamide-releasing neurons enhances appetitive behavior and food intake [32], while silencing them results in promiscuous flies [33]. The indirect inhibition from oviINs to DNc01 and DNc02 thus indicates that when oviINs are inhibited during oviposition [31], DNc01 and DNc02 can be disinhibited, putatively enhancing appetitive behavior and increasing food intake, while making flies less susceptible to mating. Correspondingly, [34] showed that females become unreceptive immediately after mating until the end of egg laying.

#### 2.2.2 Linear approach: effective connectivity with excitation/inhibition

Distinguishing excitation from inhibition is crucial for determining how the signals flow through the neural circuits. We developed a method for computing the signed effective connectivity without having to enumerate all combinations of excitation and inhibition in polysynaptic pathways. Our implementation again profitably uses a chunked sparse multiplication approach, retaining computational accessibility.

We demonstrate the capabilities of our method for computing the all-to-all signed effective connectivity for the FAFB/FlyWire connectome. We find that the effective excitation and inhibition between two cell types become similar as the path length between them increases (up to 5 hops) (Figure 1L): an analysis that would be computationally impractical without our chunked sparse matrix approach.

We computed the RFc for all cell types in the adult fly brain, considering the excitatory and inhibitory connections (Supplementary table 4). As an example, we consider here a circuit for visual looming stimulus. LPLC1 is a visual projection neuron type present in repeated instances tiling the visual field and that detects visual loom [35]. Downstream, our analysis surfaced that DNa05 integrates indirect excitatory input from LPLC1 (Figure 1M). DNa05 is a neuron type that drives flight saccades towards the ipsilateral side [36, 37]. We show further that LPLC1s provide stronger polysynaptic excitation to the contralateral DNa05, with ventral LPLC1 neurons showing stronger connections (Figure 1N). This asymmetry in the strength of the contralateral polysynaptic excitatory pathways suggests that a looming stimulus on one side would activate DNa05 more strongly on the contralateral side, causing the fly to saccade away from the stimulus. And indeed, a looming stimulus on the left activates the right DNa05, leading the fly to turn right [36]. Furthermore, the observed dorsal-ventral gradient of connectivity between LPLC1 and DNa05 (Figure 1N) may make the flies more sensitive to ventral stimuli when flying, since similar synaptic gradients are functional in directional escape responses and visual object pursuit [30, 38].

#### 2.2.3 Non-linear target neuron selection

The linear approaches described above are useful for measuring the general influence from one group of neurons to another averaged across time, but inadequate for modeling the non-linear interactions in real time, where, for the same increment in input, the neuron’s increment in activation depends on its current activation state (Figure 2A). For example, if a neuron’s activity is just below the threshold potential, then minimal additional excitation results in an action potential. To incorporate the non-linearity, we build a simple model based on the following assumptions (Figure 2B-C):

**Figure 2:**
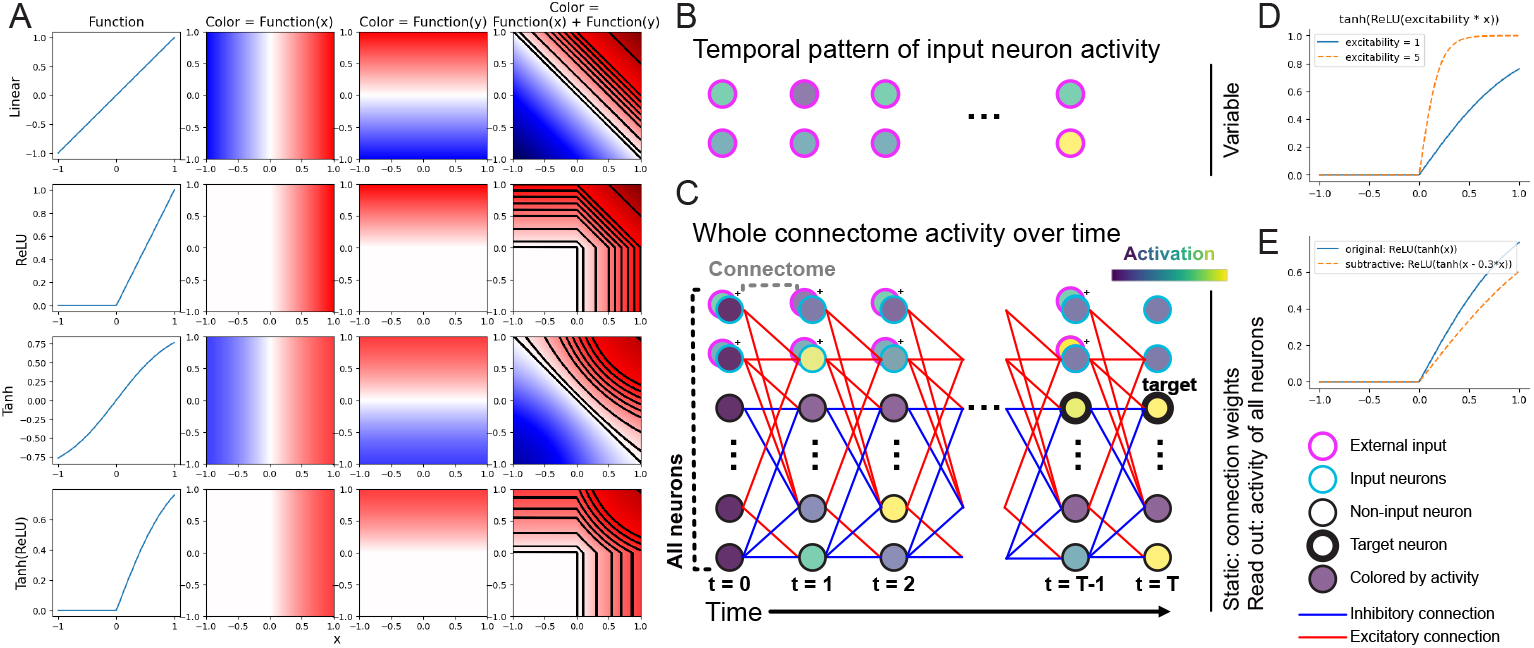
Non-linear modeling and optimal stimulus search. **A**. Non-linearity introduces context-dependence. The blue and red colors are based on the applying the function to x and y, indicated on the top of each column. Each black line in the last column indicates when the color is the same for all points on that line. When the line is straight, there is no context-dependence, i.e., the same change in x always results in the same change in color. This is the case for the linear plot, the ReLU plot when *x >* 0 and *y >* 0; and approximately true for the tanh plot when x and y are approximately 0. There is context-dependence when the lines are curved: the curvature indicates that the same change in x could result in difference changes in color, depending on the current value of x and y (the context). **B**. The input to the model is two-dimensional: the activation of input neurons across time. **C**. The model has a layered structure, where each layer represents a timepoint, and contains all neurons. The activity of all neurons is also two-dimensional (number of neurons by number of timepoints). The connectome with signed connectivity is between any two adjacent timepoints. Some neurons can be both externally stimulated (pink) and activated by upstream connectivity (blue). Neurons’ activations are illustrated with fill colors. In activation maximization, some neurons’ activation values at some timepoints are set as the target, and gradient descent looks for the two-dimensional input neurons’ activity across time that reproduces the target. **D**. The activation function of the model. **E**. Subtractive inhibition, as long as it scales with the input, is divisive in effect, i.e., it changes the slope/excitability of the target neuron.

1. A neuron cannot be ‘negatively active’, and its activation saturates;
2. A signal takes about twice the time to propagate through two synaptic hops than one;
3. Excitatory and inhibitory signals propagate at identical speeds;
4. Each neuron is modeled as a single unit.

We further require that the model be differentiable, so that experimental data can be fit via backpropagation, and for the application of activation-maximization for computing the optimal stimulus (see section 2.2.4). With these assumptions and requirements, we build a model with the following activation function:

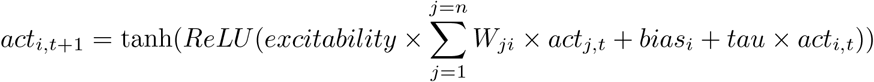

where *W*_*ji*_ (a scalar, i.e., a number) is the connection weight from neurons *j* to *i*, and *act*_*i,t*+1_ (a scalar) is the activation of neuron *i* at time point *t* +1. *bias*_*i*_ is the bias (a non-negative scalar) for neuron *i*, reflecting the baseline activity observed in experimental studies. *tau* controls the extent to which neuron *i*’s activity persists across time.

In other words, the activation of neuron *i* at the next timestep is the weighted sum of the upstream partners’ activation at the current timestep, multiplied by an *excitability* parameter (a scalar), added to the baseline activity and the remaining activity from neuron *i* itself, and then passed through a rectified linear unit (*ReLU*) and a tanh function. The *ReLU* guarantees that a neuron is not negatively active, and the *tanh* guarantees that the activation saturates. We chose tanh over a *sigmoid* for a sharper slope when the input is weak.

We further implemented the possibility of modeling a subset of inhibitory connections as contributing to divisive normalization, where inhibitory input reduces the gain of excitatory input (‘divisive’) [39], instead of subtracting from the excitatory input (‘subtractive’). Divisive normalization is implemented as a change in the *excitability* of the target neurons, scaled by the connection weight with the input, the input activation, and a user-defined divisive normalization strength parameter.

We show that the model is capable of divisive normalization without this explicit implementation (Figure 2E). For instance, suppose neuron *A* is excitatory, and neuron *B* provides feedback inhibition through *A* → *B* → *A* connections, then *the inhibition from B to A scales with A’s activity* (due to the *A* → *B* connection), rendering this inhibitory connection divisive in effect. The scaling of *B*’s activity according to *A*’s is necessary for this subtractive inhibition to be “divisive in effect”, suggesting that an explicit implementation of divisive normalization is still required otherwise.

We model temporal summation of inputs in a way that preserves differentiability, which is necessary for parameter tuning and activation maximization. In our model, a neuron only integrates presynaptic input from the timestep before. However, its presynaptic partners can accumulate activation across time through *tau*, the activity decay parameter; when *tau* is set to zero there isn’t any accummulation. When presynaptic neurons have *tau >* 0 then the postsynaptic neuron receives inputs that are representative of a temporal integration, and therefore in this way we implement temporal summation without spiking and hence maintaining differentiability.

Our model supports targeted specification of experimentally observed data by declaring simulated activity patterns, defining activation profiles over time, and customizing excitability and bias parameters, per neuron. Furthermore, our implementation supports fitting select parameters based on training data.

#### 2.2.4 Explainable-AI (XAI)-assisted interpretation

Building a model answers the question of “what would happen following a given input”, but not “what input should be given” in the first place. In the analysis of the connectome, determining which stimulus pattern best activates a select group of neurons is not straightforward given the complexity of the synaptic connectivity, which includes many divergent and convergent polysynaptic pathways with multiple lateral, feedback and recurrent connections. Furthermore, attempting to try all possible combinations of input patterns is impractical, given the high-dimensional sensory input space in the fly’s ecological niche.

We approach the interpretation of the connectome by determining optimal input patterns for select neurons anywhere in the brain with a technique from the field of Explainable AI [40]: activation maximization [41–43]. This technique manipulates inputs to optimize for the activation of target neuron(s) of interest, without searching through the entire space of possible stimuli, by formulating the search as an optimization problem that can be solved with gradient descent (Figure 2B-C). To obtain the most parsimonious stimulus and polysynaptic pathways to the target neurons possible, we define a loss function that minimizes the average activation across all neurons during the optimization.

The activation maximization approach merely delivers the optimal pattern of input neurons’ activity to reach the specified target. Therefore, the same approach supports addressing the case of setting an arbitrary, potentially time-varying, activation pattern as the target, for a select group of neurons (for example descending neurons), and then searching for the optimal activity pattern for the input neurons (for example sensory neurons).

In addition, the flexibility in specifying which neurons can be manipulated (similar to how some neurons are activated by optogenetics in a lab experiment) supports the use case of determining not only the conventional receptive field, but also the contextualised receptive field (RFc). This is useful because linking concepts, such as looming stimulus represented by LC4 [35, 44], with behaivor, can sometimes be more informative than linking individual photoreceptor activations with behaviour.

### 2.3 Pathfinding: retrieving polysynaptic pathways from inputs to targets

While the complete connectome of a whole brain can be vast, understanding of circuit function can advance by studying one subcircuit at a time. Experimental neuroscience excels at targeting select neurons in the brain, and connectomics datasets provide the basis for recovering the subcircuit between those neurons and, together with neurotransmitter signatures [10], formulating mechanistic hypotheses of circuit function.

However, identifying multi-hop polysynaptic pathways from sources to targets is computationally demanding, even impractical, in dense, highly interconnected networks like a connectome (Figure 3C). Restricting the analysis to the shortest polysynaptic pathways or thresholding graph edges to retain only the strongest connections trades off completeness for practicality, yielding potentially limited insight.

**Figure 3:**
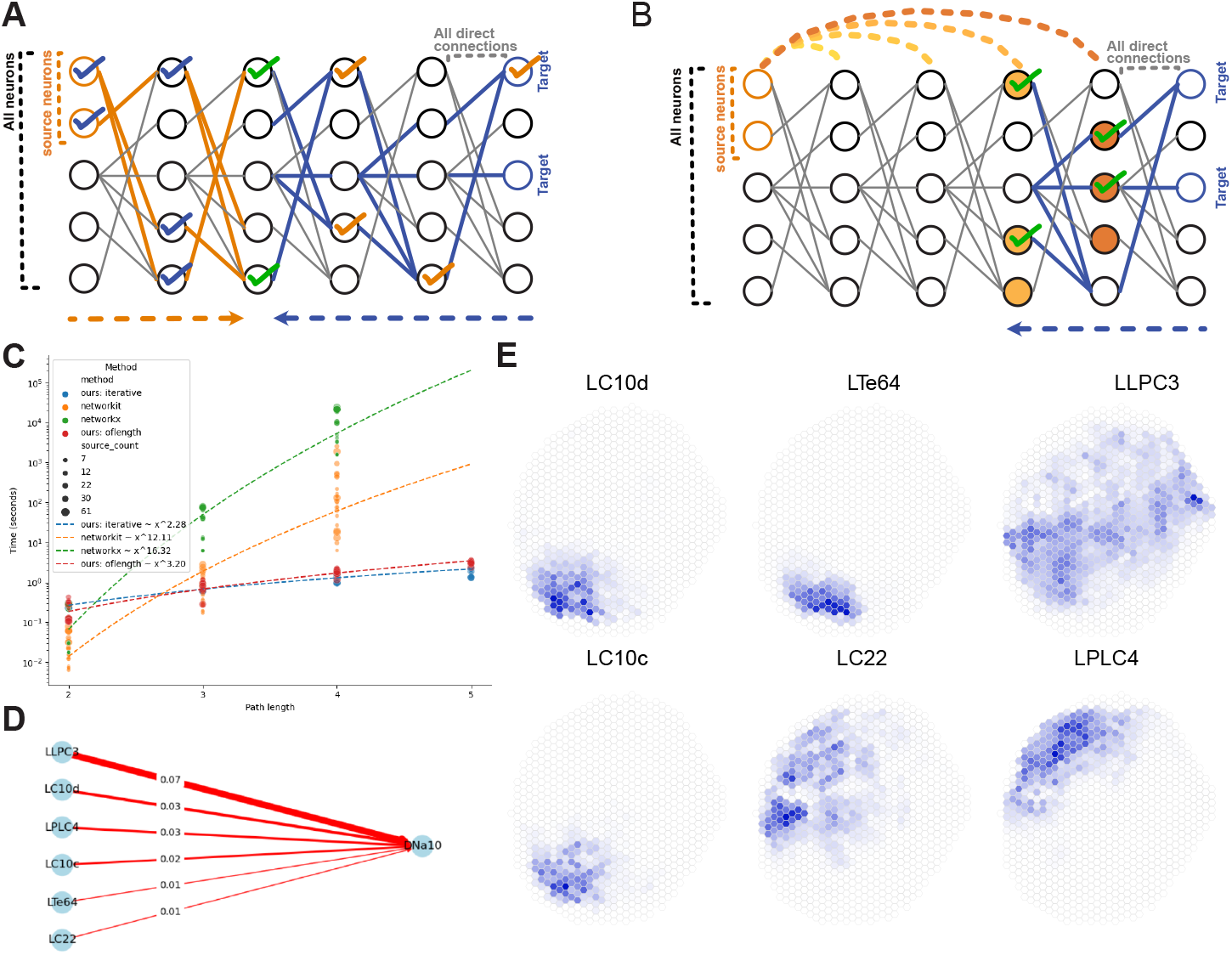
Path-finding. **A**. We identify intermediate neurons on paths of length n between source and target neurons through bidirectional breadth-first search: we propagate selection downstream from the source neurons (through the orange edges), and upstream from the target neurons (through the blue edges), until they meet at the middle, where we take the intersection of the neurons reached, those indirectly connected to both the source neurons and the target neurons (green ticks). We then propagate this selection backwards upstream (through the blue ticks) and downstream (orange ticks) to sources and targets. **B**. Alternatively, instead of propagating the selection bidirectionally, we use the pre-computed (thresholded) effective connectivity from the source (orange lines), to intersect with those connected with the targets (blue edges), starting from the last layer. **C**. We benchmark the speed of our algorithms (“oflength” refers to panel **A**, “iterative” refers to panel **B**) against Python packages networkx [45] and networkit [46], for different path lengths (x axis) and source node count. **D**. Direct connections from visual projection neurons to DNa10, where connection strength *>* 1% of DNa10’s total input. **E**. The three-hop visual receptive field through the individual neurons in each type that connects to DNa10 in **D**, shown in hexagonal heatmaps, based on the hexagonal coordinates from [47]. Equivalent three-hop and four-hop receptive fields are attached in Supplement S1.

We successfully retrieve complete circuits relating source and target neurons by introducing an efficient implementation of an algorithm that rapidly retrieves all possible circuit pathways between two neuron groups, limited only by defining the number of hops and an optional threshold on the connection weight. Our approach yields results in seconds for connectomes such as the adult fruit fly brain in consumer-grade laptops (Figure 3C).

#### 2.3.1 An efficient search algorithm

First we represent the connectome as a table (a Python Pandas dataframe with indexed columns) with *n* rows, where each row is a connection with 3 properties: the presynaptic neuron ID, the postsynaptic neuron ID, and the connection weight, with an O(*n*) cost. Second, perform a bidirectional breadth-first search, by subsetting the table once per hop in each direction, using the set of IDs of the neurons of each group (source and target) as keys to retain, and proceed iteratively. The termination condition is reaching half the specified number of hops (or half minus one for one end when odd) from each side. The result is the complete set of edges or connections, relating all neurons in the circuit between the source and target neurons and spanning the specified polysynaptic pathway depth.

We also implemented a second method, consisting in propagating the neuron selection upstream from the target neurons only, and intersect those connected to the target neurons with those indirectly connected to the source neurons, using the precomputed effective connectivity (Figure 3B). Though slightly faster, this algorithm relies on the (thresholded) effective connectivity matrices, which by design (for performance tuning and dramatic data storage savings) can be constructed to leave out weak connections via thresholding.

Both methods retrieve the synaptic graph of the subcircuit of interest relating all source and target neurons. If the complete list of polysynaptic pathways was required (where each pathway contains *n* edges from a source to a target for a path of length *n*), depth-first search can proceed based on the extracted subcircuit only, so that every run returns a hit, bypassing the unsuccessful searches therefore gaining in performance.

Our approach significantly reduces unnecessary operations, enabling interactive path-finding in mere seconds on widely accessible consumer hardware or Google Colab, improving performance substantially over conventional methods that can take hours.

#### 2.3.2 Example search: a visual receptive field

In the fruit fly brain, the optic lobe alone contains about 50,000 neurons on one side [9]. The circuits for visual processing are organised in repeated columnar units corresponding to ommatidia that tile the surface of the eye, capturing a two-dimensional slice of the visual field in a retinotopic map [9, 48, 49]. Deeper in the brain, neural circuits for input convergence across the visual field results in individual neurons responding to specific visual features [21].

To showcase our approach to swiftly retrieve neural circuits between source and target neurons separated by polysynaptic pathways, we investigate the visual receptive field of DNa10, a descending neuron cell type hypothesized to control wing thrust, lift, and pitch on the basis of connectomics data [37]. Our preparatory analysis revealed that DNa10 receives over 1% of its direct synaptic input from visual projection neuron types LLPC3, LPLC4, LC10d, LC10c, LTe64, LC22, each of which is composed of individual neurons tiling the visual field (Figure 3D).

Assuming different VPNs detect different visual features, we ask whether the visual features DNa10 detects are location-dependent. We retrieved 3-hop polysynaptic pathways, from the large monopolar cells *L*1, *L*2 and *L*3 (directly postsynaptic to photoreceptors), and photoreceptors *R*7 and *R*8, to DNa10, separately through each VPN type. Based on the columnar assignments of the large monopolar cells and photoreceptors in [47], we group the pathways by hexagonal coordinates tiling the visual field, and calculate the effective connectivity from these coordinates to DNa10, to generate the hexagonal heatmaps describing the visual-feature-specific receptive field (Figure 3E). The equivalent plots for 4-hop and 5-hop visualizations are in Supplement S1. The visual features that DNa10 can respond to could thus vary depending upon the area of the visual field, which is plausible for a descending neuron involved in flight control.

#### 2.3.3 Structuring circuits into layers

Structuring a circuit into layers facilitates the identification of motifs of known computational properties such as feedforward, feedback and lateral inhibition, as well as winner-take-all architectures and recurrent motifs where prior processed signals interact with newly incoming signals. To this end, we integrated a metric of information flow [50] which labels neurons into layers as a probabilistic function of the connection weight (see Methods 5.1).

### 2.4 Depth and density of brain circuits

With the ability to swiftly retrieve polysynaptic pathways across the whole fly brain connectome, we set to measure how many hops separate any one cell type from another. We sample 100 random cell types as sources, and another 100 random cell types as targets, and then search for polysynaptic pathways from 1 single source cell type to 1 single target cell type at a time. Across all 10,000 possible pairs of cell types, we found that, for a connection threshold of 0, ∼ 70% of tested pairs yielded pathway lengths of 2 (i.e., one intermediate neuron in between), a result that changes dramatically for a connection threshold of 1%, yielding only 2% of the pairs with a pathway of 2 hops (Figure 4A). For polysynaptic pathways of 5 hops, the difference is much smaller between a threshold of 0 (100%) and 1% (∼ 84%). These findings indicate that the fly connectome is very dense when considering weak connections or polysynaptic pathways of longer length, with significant implications for reductionist approaches to studying brain function.

**Figure 4:**
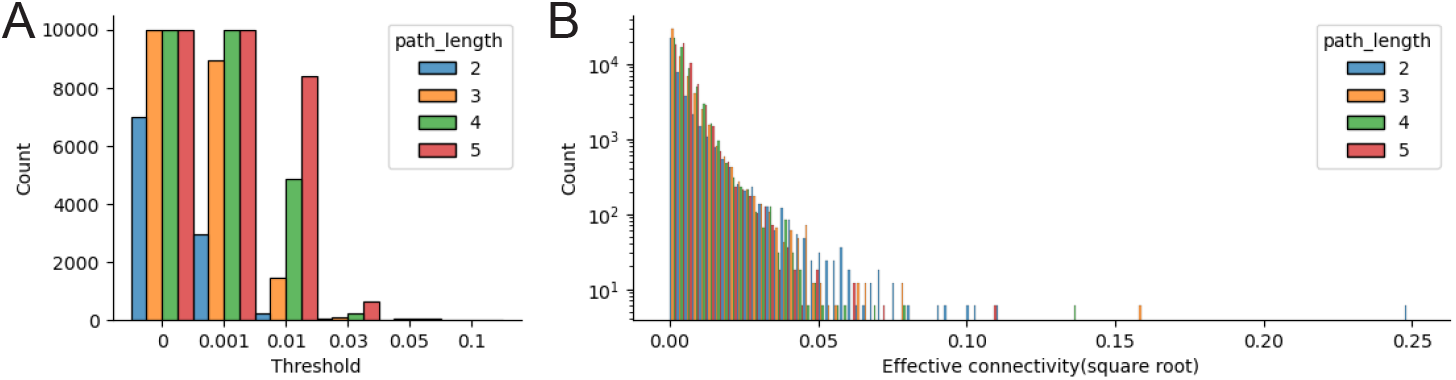
Density and effective connectivity of polysynaptic pathways. We randomly select 100 cell types as sources, and 100 as targets from the FAFB/FlyWire connectome. We look for poly-synaptic pathways between them one pair at a time, and calculate effective connectivity when such pathways exist, for all 10,000 pairs, for path lengths between 2 and 5. **A**. Histogram of the number of cell type pairs with paths of the tested lengths, given a threshold on all the direct connections in the pathways. **B**. Histogram of effective connectivity between source and target cell types for the paths found. The x axis are square-root-transformed, and the y axis log-transformed for visualization.

### 2.5 Impact of circuit density on effective connectivity

Using the pathways between the selected cell types, we compute the effective connectivity locally, to compare the effective connectivity value across path lengths. Effective connectivity calculation typically uses connection weights normalized by the total number of post-synapses for the postsynaptic neuron, and is therefore a number between 0 and 1. Taking the product of such numbers intuitively results in a smaller number the more times the multiplication takes place. However, we show here that effective connectivity is similar in magnitude across path lengths. This is likely related to the density of the connectivity, i.e., the abundant divergence and convergence in the nervous system, such that there are more pathways between two cell types the longer the path length is. The accumulation of these pathways likely counteracts the diminishing product of connection weights.

### 2.6 Effect of non-linearity

Non-linearity introduces context dependence with important consequences for neuronal activation when integrating different types (e.g., excitatory or inhibitory) of upstream signals with different weights. Here, we consider two toy circuits to exemplify when a non-linear model produces neural dynamics different from a linear model. The first example illustrates the role of *tanh* (saturation) and the second the role of *ReLU* (non-negativity).

#### 2.6.1 Modeling convergent feedforward excitation and inhibition to explore the role of *tanh*

In the first example, we explore how the difference in weights for feedforward excitation and feedforward inhibition changes the activity of the output neuron across varying strengths of input.

To demonstrate the differences between the linear and non-linear neuron activation responses to increasing inputs, we constructed the following network model with parallel feedforward excitation and feedforward inhibition converging onto a single output neuron (Figure 5A). Hence, a single input unit transmits signals to two units (the interneurons), one excitatory, with a stronger weight, and one inhibitory, with a weaker weight, and both are connected to the single output neuron. The strength of the weights (*w*1 and *w*2) represent here the aggregate convergent input from many neurons, which can co-activate in ecologically-realistic environments. We model this convergence with one unit for simplicity.

**Figure 5:**
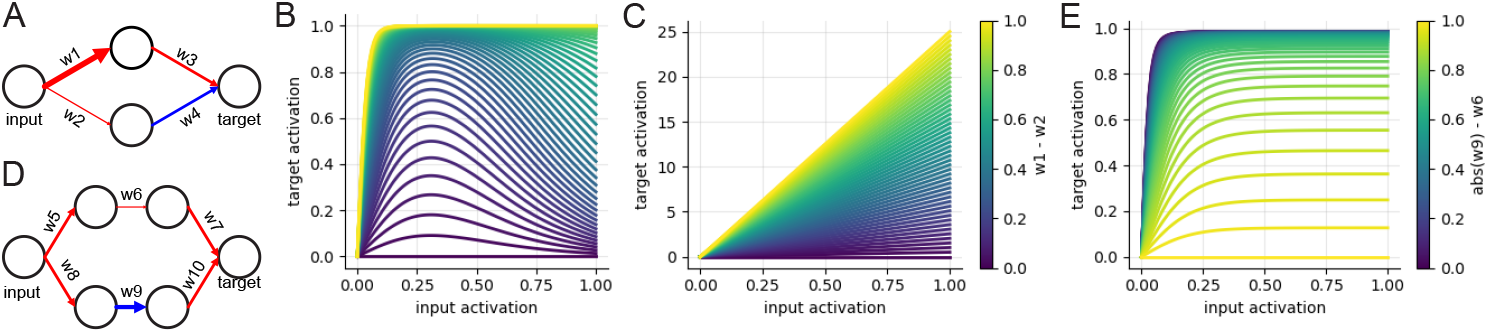
Toy circuits for probing non-linearity. **A**. A simple circuit with both feed-forward excitation (*w*1 → *w*3) and feed-forward inhibition (*w*2 → *w*4). We assume *w*3 = *w*4. **B**. Higher input activation may not lead to higher target activation in a non-linear model, depending on the difference in saturation (here due to the difference between *w*1 and *w*2) between feed-forward excitation and feed-forward inhibition. **C**. Higher input activation leads to higher target activation in a linear model, as long as effective excitation *>* effective inhibition. **D**. A simple circuit where the effective inhibition (through *w*8 → *w*9 → *w*10) outweighs the effective excitation (through *w*5 → *w*6 → *w*7). However, activation of the inputs in the non-linear model activates, instead of inhibits, the target (**E**.), because the neuron downstream of *w*9 is silenced and does not pass on the inhibition.

With the non-linear model, we find that the output neuron’s activity is not always the highest when the input neuron’s activity is the highest (Figure 5B). In contrast, higher input entails higher output for the linear model (Figure 5C). Our reasoning is the following. The up-and-down pattern of output neuron responses in Figure 5B arises from the differences in the excitatory drive onto the excitatory (*w*1) and inhibitory (*w*2) interneurons. When *w*1 *> w*2, at a high input value, the excitatory interneuron saturates first. At even higher inputs, the excitatory interneuron can no longer increase the excitatory drive to the target, therefore limiting the feedforward excitation, whereas the inhibitory drive can be higher still. This effect stems from the saturating non-linear region of the *tanh* function (Figure 2D), supporting its use in our neuron model.

We further explored this phenomenon by varying the difference in connection weights from the input neuron onto the excitatory (*w*1) and inhibitory (*w*2) interneurons (when *w*1 + *w*2 = 1) (Figure 5B). When *w*1 = *w*2 = 0.5, the excitation and inhibition onto the target neuron cancel out; when *w*1 − *w*2 = 1, *w*2 = 0, the output neuron only receives excitation, so it quickly saturates; the nonlinearity in response manifests the most when 1 *> w*1 *> w*2 *>* 0, where the excitatory interneuron approaches saturation, and the inhibitory interneuron, while providing inhibition, does not approach saturation yet.

This example highlights the context-dependence in non-linear models: the excitatory interneuron is approaching saturation, so further excitation makes little difference in its activation; in contrast, the inhibitory interneuron does not saturate yet, so little excitation leads to substantial increase in activity. Together, they cause the non-linear response in the output neuron.

#### 2.6.2 Differential transmission of inhibition and excitation: role of the non-negativity constraint with *ReLU*

In the second example, we explore how the existence of inhibitory neurons, while enforcing a non-negativity constraint on neuron activity, favors the use of a non-linear model over a linear one. Here, the non-linearity of *ReLU* plays a key role by introducing a non-negativity constraint: when an interneuron receives more inhibition than excitation (such as that in Figure 5D), the net activity is zero, instead of negative, and therefore does not pass on any of such excess inhibition – just like in biological neurons. The target neuron thus receives less inhibition than the connection weight alone would suggest if a linear model was used.

Together, these two examples highlight how simple non-linearities - saturation via *tanh* and thresholding via *ReLU* - can reshape circuit responses in ways not predictable from linear connectivity alone.

## 3 Discussion

We have introduced the Connectome Interpreter (https://github.com/YijieYin/connectome_interpreter), an open-source computational toolbox that translates the complex structural wiring of a brain into testable functional hypotheses. We demonstrated that out approach for computing effective connectivity, with or without excitation/inhibition, scales efficiently to large connectomic datasets such as the adult fruit fly brain with ∼ 140,000 neurons and millions of synaptic connections [2, 3] with predicted neurotransmitter signatures [10]. By integrating the fly connectome with a curated and community-expandable compendium of known neuronal functions, our methods move beyond a static anatomical map to enable the dynamic exploration of signal propagation, both linearly and non-linearly, across the entire brain. Our toolkit facilitates combining experimentally acquired neuronal activity data with the connectome and the synaptic signatures to infer *in silico* the receptive field of any neuron in the brain, and to slice out relevant subnetworks between select sets of source and target neurons, aiding in bridging the gap between circuit architecture and neural computation.

### 3.1 The need for the whole connectome to analyze individual neurons and circuits

The fruit fly connectome is very dense (Figure 4A; [51]). By this we mean that the activity of any one neuron will depend on the activity of many other neurons. Therefore, our intuition falls short in formulating hypotheses of when or whether a neuron would be active as a function of the activity of some other neurons.

With our toolkit, we measured the density of 5-hop polysynaptic connectivity between random cell types, at 1% threshold for normalized input and not considering the sign of the connections. We found that for 84% of all possible pairs of cell types there was at least 1 polysynaptic path (Figure 4A), which strikes us as very high. Our finding highlights the abundance of convergent and divergent connections in the nervous system of the fly, and has implications in the analysis of circuit function.

For example, let’s consider action selection in the fruit fly. The observed dense connectivity fully supports how the fruit fly, in response to a specific sensory stimulus, selects one course of action while suppressing others, because all other inputs must be integrated towards taking the decision, which requires strong convergence. Furthermore, all other possible courses of action have to be suppressed, which requires both divergence (lateral inhibition) and feedback inhibition to setup expectations via corollary discharge. With the Connectome Interpreter toolkit, dense connectomes can be analyzed with ease to assist in formulating hypotheses of neural circuit function that can be tested experimentally in the laboratory by monitoring and manipulating neuronal activity.

### 3.2 Necessity of non-linear models

Biological neurons are non-linear in nature, given a dynamically adjustable temporal integration window, a spike that resets input accumulation and transmits signals to postsynaptic neurons, and a refractory period that limits maximal activity, by limiting the maximum number of spikes per unit of time (the firing rate). A consequence of spiking is that neurons are either silent (zero) or active (positive), and cannot be negatively active. While graded potential neurons exist, for example in the optic lobe of the fly [52], their activity dynamics is also constrained to be non-negative and are subject to thresholds for synaptic transmission, both of which are better represented by non-linear functions.

A good model captures enough from the system under study that insight can be derived from the study of the model that would not be possible otherwise, often because of the impracticality of observing the whole system with enough spatial and temporal resolution. What makes a good model then depends very much on the goals of the desired analysis. Here, we set to capture fundamental features of neuronal activity such as the non-negativity constraint and the saturation of activity, features that all reflect different aspects of a neuron’s non-linear activation function, while retaining differentiability for using experimental data to fit model parameters and for computing the optimal stimulus with activation maximization.

We modeled the saturation of neural activity with a *tanh* and the non-negativity constraint with a *ReLu* centered at zero. In biological neurons, *tanh*-like saturation has been shown experimentally [53], and the prevalence of such saturation in neural responses likely depends on the overall activity level of the system: as network activity increases, more neurons approach their firing-rate ceiling and saturate. The *ReLU* function trivially implements the non-negativity constraint by discarding all inputs below a defined threshold, here being zero. The non-negativity constraint models an effective deficit in inhibitory transmission: the fact that inhibition does not propagate beyond the postsynaptic neuron as negative signals would when not prevented to do so. The effective deficit in inhibitory transmission requires only that a neuron receives more net inhibition than the amount that would completely silence it by canceling out excitatory input. Given that essentially all brain circuits are composed of both excitatory and inhibitory neurons, the condition of effective inhibitory deficit is presumably widespread, indicating that the non-negativity is fundamental to capture in the model for the propagation of excitatory and inhibitory signals across the connectome.

We tested our model and found that nonlinearities such as saturation and thresholding can have non-intuitive effects, where higher input activation may not entail higher target activation (Figure 5A-B), due to *tanh*-based saturation of neurons; and where effective net inhibition may not entail the inhibition of the target (Figure 5D-E), due to the silencing of the inhibitory path through *ReLU*.

### 3.3 The Connectome Interpreter toolkit in practice

We show the utility of combining existing literature on neuron function with effective connectivity, by highlighting previously unreported links between neurons with known functions (Figure 1H-K, Figure 1M-N, Figure 3D-E). The interpretability of these links underscores the promise of contextualized receptive fields (RFc) for hypothesizing on the functions of currently understudied neurons. Together, effective connectivity and contextualized receptive field provide a principled way to move beyond direct synapses, link indirectly-connected neurons representing high-level concepts, discover putative functional relationships between neurons, and generate testable hypotheses of neural function.

Our model provides flexible capabilities to incorporate experimentally-tested features of the nervous system, such as divisive normalization, baseline activity, excitability, and persistence of activity across time. The interpretability of the parameters makes incorporating and testing assumptions straightforward, where the model serves as a conduit for propagating activity, and a hypothesis-generation tool by answering “what-if” scenarios under plausible assumptions. The library of functions we provide facilitates the interpretation and visualization of the model results, guiding the analysis of the model’s predictions.

The toolkit includes library utility functions for calculating effective connectivity and path-finding, that operate orders of magnitude faster than other methods. The performance improvement we achieved, together with the accessibility afforded by lowering the minimum hardware requirements, democratizes data analysis and therefore supports a qualitative change in approaches to connectomic analysis. We offer a range of examples in the form of annotated programs as online notebooks, alongside detailed documentation, for easy adoption. The code for reproducing much of the analyses in this manuscript is also available in this Google Colab notebook. Our software, implemented as open source and thoroughly documented, can be used as a standalone or incorporated into existing data-query platforms such as neuPrint [54], Codex [20] and CAT-MAID [55], to further lower the barriers of access and facilitate the discovery of neurons and circuits relevant to any and all experimental paradigms inquiring into brain function.

## 4 Acknowledgements

We thank Mateo Espinosa Zarlenga for his helpful advice on activation maximization. We thank Laura Burnett and Arthur Zhao for their contributions and support, especially on the incorporation of hexagonal and anatomical receptive field plotting capabilities, and for initial testing of the information flow algorithm. We thank Michael Reiser for facilitation and support of collaborations. We thank Alexander Shakeel Bates, Sophia Renauld, Diego A. Pacheco, Mathew F. Collie, Helen H. Yang, Jingxuan Fan and Arie Matsliah for early access to cell function annotations related to reconstructions in the BANC, MANC and FAFB connectomes, now released with the BANC paper, as well as additional inferences from published work. We thank Michael Reiser, Judith Hoeller, Xin Zhong, and Fred Wolf for contributions to the table of neurons with known functions. We thank Katherine Nagel, Kavin Nuñez and Shamik Dasgupta for their feedback and suggestions on the package. We thank Gregory S.X.E. Jefferis for his support. We thank Arie Matsliah for incorporation of the table of neurons with known functions into Codex. We thank the LMB High Performance Computing facilities for the computational resources. This project was funded by a Wellcome Trust Investigator Award to Albert Cardona (Ref: 205038/Z/16/Z) and the MRC LMB core funding.

## 5 Methods

### 5.0.1 Calculating effective connectivity

A standard approach is to multiply the weights in a multi-hop path between two neurons, and add the products across parallel pathways [16, 18, 19, 56], without any non-linearity such as thresholding or saturation. This can be implemented as multiplication of the connectivity matrix with itself (see explained in Figure 1C). Calculating matrix power scales to the third power of the number of nodes because: for matrix multiplication *A* × *B* = *C*, to calculate each element in *C, n* multiplications need to take place (*n* elements in a row of *A*, each multiplied with the corresponding element in a column of *B*). There are *n* × *n* elements in *C*.

While theoretically straightforward, this approach rapidly encounters practical limits: Storing a 140,000 x 140,000 connectivity matrix in computer random access memory (RAM), where each element takes 32 bits (float32), requires:

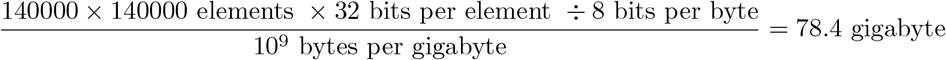

, exceeding the capacity of common laptops. Matrix multiplication requires at least three times this amount (result = one matrix x another matrix). Matrix multiplication is faster on GPUs than CPUs (Figure 1F). However, the maximum RAM an A100 chip has is 80GB, indicating that additional parallelization is necessary, even on high performance computing clusters, to take powers of the dense matrix the size of an adult fly brain.

One inherent aspect of calculating polysynaptic strengths by multiplication is the rapid diminishing (for normalized weights) or amplification (for synapse counts) of values as the pathway length increases. We believe this accurately reflects divergence and convergence in the nervous system: the signal in one neuron can be traced to many in a small number of hops, corresponding to the increase in connection density (Figure 4A). Our default option therefore does not process the products further.

Nonetheless, users have the flexibility to normalize these pathway strengths using an n-th root transformation, to “control for the path lengths”. However, due to the divergence and convergence, the rooted effective connectivity values tend to be bigger with longer path lengths. This limitation of diminishing or amplification of values is better addressed using the non-linear model (Section 2.2.3).

### 5.1 Inferring Hierarchy from Connectivity to Sensory Neurons

To facilitate interpretation of large connectivity matrices, we define a hierarchy of neurons roughly based on how many hops away a neuron is from sensory neurons. This hierarchy approximates the feedforward processing of information from sensory neurons toward (motor-proxy) descending neurons, while revealing deviations due to lateral and feedback connectivity.

In an idealized feedforward network, neurons are organized into discrete layers based on the number of hops that separate them from inputs. However, neurons in real brains are more richly interconnected. For example, a sensory neuron may strongly connect to neuron A and only weakly connect to neuron B, and A strongly connects to B. In this case, we have to decide if neuron B is one hop or two hops away from sensory neurons. An additional complication is that downstream neurons may connect back to sensory neurons, obfuscating a simple layering.

We adopt a nonlinear signal propagation model previously developed for the Drosophila olfactory system [50], which simulates the hierarchical spread of activity from an initial set of active neurons. The model treats each neuron as binary–active or inactive–and updates activity in discrete steps. At each step, inactive neurons are probabilistically activated based on their total weighted input from already active neurons. This probability follows a sigmoid function, capturing both integration and saturation. Once active, a neuron remains active for all subsequent steps.

The layer of a neuron is defined as the average number of steps it takes to become active in this simulation, given a starting set of input neurons. These layers capture a continuous (non-integer-valued) position in a hierarchy that starts at the input neurons. The model is implemented using the Navis toolbox [57] and allows users to visualize pathways together with the hierarchical position of neurons. The hierarchy can also be used to identify feedforward, lateral, and feedback neurons, i.e., neurons which predominantly connect to other neurons of later, similar, or earlier layers, respectively.

### 5.2 Modeling

For user friendliness, we implement parameter-sharing: for parameters such as bias, slope, and divisive normalization, users need to first specify how the neurons are to be groups (by e.g. cell type), and then simply use dictionaries that map from the group names to the values of the corresponding group.

We also provide training capability. Given that data from the experimental literature are not always time series, but sometimes a number representing average activity across time, we support training using either the average activation value for each neuron, or time series data. Parameter sharing also applies to training, such that each neuron group have the same parameter values after training.

#### 5.2.1 Saliency

We implement another technique from explainable AI, saliency, which calculates the gradients of the target with respect to the input. Compared to activation maximization, which looks for the best stimulus to reach the target activation, saliency addresses the question of “which part of the current stimulus (i.e. which input neurons, among all activated) matters more for the current target neuron activation?”. Note that due to non-linearity, the importance only refers to the current activation value for both the input and the target. The important part of the stimulus may change for a different target neuron activation value.

### 5.3 User-centric design

We aim for user-centric design in writing the package, and therefore included a number of convenience functions for visualization of pathways in (interactive) wiring diagrams, with options to sort nodes by connection weight, or user-specified order. Effective connectivity can not only be visualized as colored tables, but also in neuroglancer, or for visual information, in hexagonal heatmaps or anatomical receptive fields [58]. Pathways can also be plotted through layers in neuroglancer.

To make the non-linear model interpretable, we implement a function that extracts the “relevant neurons” - those that are connected (poly-synaptically) to the target with a connectivity and activity threshold. Conversely, users can start from a pathway and query the activity of all neurons in a pathway. In addition, the activity evolution of groups of neurons can be visualized in an interactive plot, or in neuroglancer.

To facilitate research on vision and navigation, we add functions that construct looming stimuli and sine-shaped stimuli.

To further reduce the barrier of linking stimulus space to neuron activation space, we collate published datasets of neruon responses to odour presentations [53, 59**?** –62], so that the community can easily combine experimental literature with connectome-based modeling.

Last but not least, we illustrate the functionalities of the software through example Google Colab notebooks (https://github.com/YijieYin/connectome_interpreter), using public datasets, to enable browser-based analysis without local installation.

## 6 Supplementary materials

**Figure S1:**
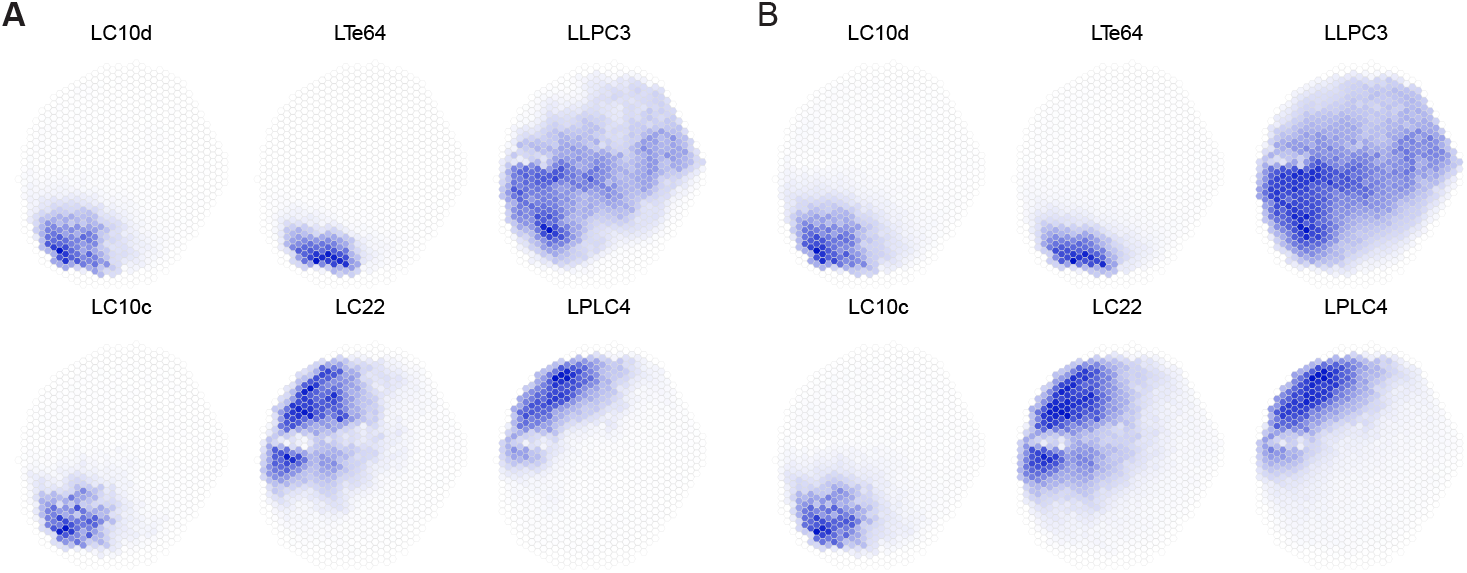
Four-hop (**A**) and five-hop (**B**) visual receptive field through the individual neurons in each visual projection type that connects to DNa10 in Figure 3D

1. Supplementary table 1. A snapshot of the table of neurons with known functions.
2. Supplementary table 2. Direct connectivity combined with table of neurons with known functions.
3. Supplementary table 3. Effective connectivity combined with table of neurons with known functions.
4. Supplementary table 4. Signed effective connectivity combined with table of neurons with known functions.

